# A Rare Constitutional Centromeric Expansion of Chromosome 1 in a Phenotypically Normal Individual: A Comprehensive Cytogenetic Study

**DOI:** 10.64898/2026.11.12.766333

**Authors:** Kapil B. Shirsath, Utkarsha N. Desai, Aarti Notnani, Rajesh Kumar Chaurasia, Nagesh N. Bhat, Arshad Khan

## Abstract

Constitutional centromeric heteromorphisms are uncommon chromosomal variants that may complicate cytogenetic interpretation. We report a constitutional enlargement of the centromeric region of one chromosome 1 homologue identified incidentally during biological dose estimation in a suspected occupationally exposed individual. Conventional cytogenetics, pan-centromeric fluorescence in situ hybridization (FISH), and multiplex FISH confirmed the variant and excluded cryptic chromosomal rearrangements. Analysis of 500 metaphases revealed no dicentric chromosomes. Quantitative assessment demonstrated a centromere-to-chromosome length ratio of 0.314 ± 0.019 for the enlarged homologue compared with 0.169 ± 0.019 for the unaffected homologue and 0.161–0.186 in controls, representing an approximately 1.86-fold increase (t = 28.4, p < 0.0001). The findings establish a benign constitutional centromeric heteromorphism of chromosome 1 and highlight the importance of distinguishing such variants from structural chromosomal abnormalities during routine cytogenetic evaluation.

## Introduction

Centromeres are essential chromosomal domains that ensure accurate chromosome segregation during mitosis and meiosis, thereby maintaining genomic stability. Structural or functional alterations affecting centromeres can lead to chromosome mis-segregation, aneuploidy, and diverse pathological outcomes (1, 2). In humans, centromeres are primarily composed of large arrays of highly repetitive α-satellite DNA organized into chromosome-specific higher-order repeat (HOR) structures (3, 4). Considerable variation exists within these arrays, including differences in sequence composition, HOR organization, and overall array size between homologous chromosomes and among individuals (3-5). Such variation contributes to constitutional chromosomal heteromorphisms, which are generally regarded as benign polymorphic variants lacking explicit phenotypic consequences (6). Chromosome 1 is particularly notable for its large and polymorphic centromeric and pericentromeric repetitive DNA content, making it one of the most structurally variable regions of the human genome (3, 5). Although heteromorphic variants involving the pericentromeric heterochromatic block of chromosome 1 (1qh variants) are relatively well recognized in routine cytogenetics, alterations involving the centromere itself have been reported far less frequently and remain incompletely characterized (6). Despite advances in fluorescence in situ hybridization (FISH) and molecular cytogenetics, most published descriptions of constitutional centromeric variants rely predominantly on qualitative morphological assessment. Quantitative evaluation of centromeric enlargement and comprehensive confirmation using complementary cytogenetic approaches remain uncommon. Consequently, the prevalence, extent, and cytogenetic characteristics of constitutional centromeric expansions are likely underreported, particularly when identified incidentally during routine chromosome analysis. This knowledge gap is relevant because unusual centromeric morphologies may be mistaken for structural chromosome abnormalities, potentially complicating cytogenetic interpretation in clinical genetics, cancer cytogenetics, and radiation biodosimetry investigations. Accurate characterization of such variants is therefore important for distinguishing benign constitutional polymorphisms from clinically significant chromosomal abnormalities.

Here, we report a constitutional enlargement of the centromeric region of one chromosome 1 homologue identified incidentally during biological dosimetry analysis and provide detailed cytogenetic characterization of the variant using conventional karyotyping, pan-centromeric FISH (pan-cent-FISH), and multiplex FISH (mFISH).

## Materials and Methods

### Chemicals

ProLong™ Diamond Antifade Mountant with DAPI was obtained from Invitrogen (Thermo Fisher Scientific, USA). Giemsa stain, DPX mountant, streptomycin, and penicillin were sourced from Sigma-Aldrich (USA), Merck (USA), and HiMedia (India), respectively. RPMI 1640, phytohemagglutinin (PHA), fetal calf serum (FCS), and colcemid were procured from Gibco Life Technologies (USA). Pan-centromere and mFISH probes were obtained from MetaSystems (Germany).

### Ethical Approval and Blood Collection

The study was approved by the Institutional Ethics Committee of Bhabha Atomic Research Centre, Mumbai, India (Ref: BHMEC/NP/12/59/2026). Peripheral blood (4 mL) was collected from two male donors; one subject and one healthy volunteer, both without relevant medical history. Written informed consent was obtained from all participants, and all procedures adhered to established ethical guidelines.

### Blood Culturing, Metaphase Harvesting, and Giemsa Staining

Sixteen whole blood cultures (eight per donor) were established following our previously optimized protocol (7), in accordance with IAEA (8) and ISO (9) guidelines, using RPMI-1640 supplemented with FCS, PHA, L-glutamine, and antibiotics. Colcemid (100 µL; 250 µg/mL) was added at 24 h to arrest cells in metaphase, and chromosomes were harvested at 52 h. Cells underwent hypotonic treatment and fixation in Carnoy’s fixative, after which fixed suspensions were dropped onto clean slides to prepare metaphase spreads. Slides were treated with RNase and stained with Giemsa for cytogenetic analysis.

### Pan-Centromeric FISH

Two slides were hybridized with fluorescently labelled pan-centromere FISH probes following standard protocols (10). Briefly, slides were subjected to pepsin treatment, after which 16 µL of probe mixture was applied, denatured at 75°C for 3 min, and hybridized overnight at 37°C in a humidified chamber. Post-hybridization washes were performed in 0.4× SSC at 72°C, followed by a brief rinse in 2× SSC/0.05% Tween-20. Slides were dehydrated through a graded ethanol series (80%, 90%, 100%) and mounted in DAPI-containing antifade mountant.

### Multiplex FISH (mFISH)

Two slides were processed for mFISH per standard protocols (11, 12). Briefly, slides were pepsin-treated (1 g/50 mL, 0.01 N HCl; 37°C, 2 min), post-fixed in 1% formaldehyde/1× PBS/50 mM MgCl□ (10 min, RT), denatured in 0.07 N NaOH (1 min, RT) following a 2× SSC incubation (70°C, 30 min), and dehydrated through a graded ethanol series. Denatured probe (75°C, 5 min) was hybridized over a 22×44 mm^2^ area for 3 days at 37°C. Post-hybridization washes (1× SSC, 75°C; 4× SSCT; distilled water) were followed by air-drying and mounting in DAPI-containing antifade mountant for microscopic examination.

### Imaging and Analysis of Metaphases

Slides were scanned using the Metafer5 system (MetaSystems, Germany) on a ZEISS Axio Imager.Z2 equipped with a motorized stage and CoolCube 4 CCD camera, with metaphase detection at 10× and image capture at 63× (AutoCapt mode). Giemsa and FISH slides were analyzed using the Ikaros and ISIS modules, respectively (7). Pan-centromere FISH images were captured under FITC and DAPI filter sets, while mFISH metaphases were sequentially acquired under six filter sets (FITC, Spectrum Orange, Texas Red, Cy5, DEAC, and DAPI) and overlaid for chromosome identification.

### Statistical Analysis

The centromere-to-chromosome length ratio (CCR) was calculated for a minimum of 50 metaphases per donor, with results expressed as mean ± SD. Inter-individual differences were assessed using Student’s unpaired t-test (p < 0.05).

## Results

### Case History

Cytogenetic analysis of a peripheral blood sample from a suspected occupationally exposed individual was conducted for biological dose estimation at the Biodosimetry Laboratory, Bhabha Atomic Research Centre (BARC), Mumbai, India. Scoring of 500 metaphases revealed no dicentric chromosomes, indicating no evidence of significant radiation exposure.

During routine chromosomal analysis, a constitutional centromeric abnormality was incidentally identified on one homologue of chromosome 1, present consistently across all metaphases examined, while the other homologue appeared cytogenetically normal (Figure 1). The variant was classified as constitutional given its presence in every metaphase analysed. The individual displayed no associated phenotypic abnormalities, and available family history indicated that both parents and offspring were phenotypically normal.

**Figure 1:**
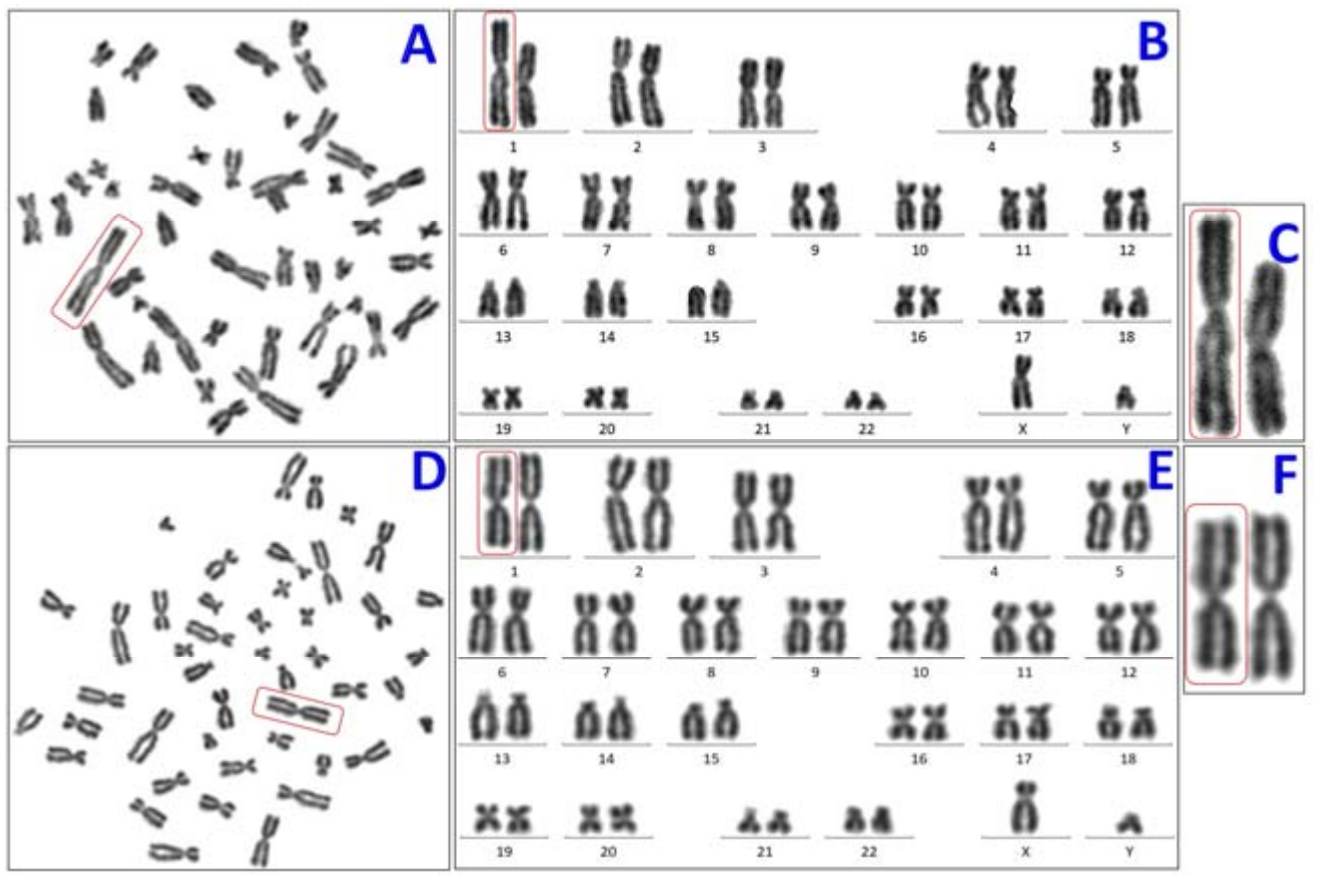
Representative Giemsa-stained metaphase chromosome preparations. (A–C) Metaphase from a subject exhibiting an enlarged centromere in one homolog of chromosome pair 1. (A) Representative metaphase spread. (B) Corresponding karyotype. (C) Enlarged view of chromosome pair 1 highlighting the enlarged centromere (red box). (D–F) Representative metaphase from a healthy volunteer (control). (D) Metaphase spread. (E) Corresponding karyotype. (F) Enlarged view of chromosome pair 1 showing normal centromere morphology. Red boxes indicate the chromosome pair selected for detailed visualization.

### Centromeric Heteromorphism of Chromosome 1

Giemsa-stained metaphase spreads from the subject and an age and sex-matched healthy control were initially examined to assess centromeric morphology. Conventional cytogenetic analysis revealed a marked enlargement of the centromeric region of one chromosome 1 homologue in the subject, whereas the second homologue and both chromosome 1 homologues in the control displayed typical centromeric morphology (Figure 1).

To further characterize and validate this observation at higher resolution, metaphase preparations from both individuals were analysed using pan-centromeric fluorescence in situ hybridization (pan-cent-FISH) (Figure 2) and multiplex FISH (mFISH) (Figure 3). Both techniques independently and consistently confirmed the presence of an abnormally expanded centromeric region restricted to one chromosome 1 homologue in the subject, with no comparable finding in the control.

**Figure 2.**
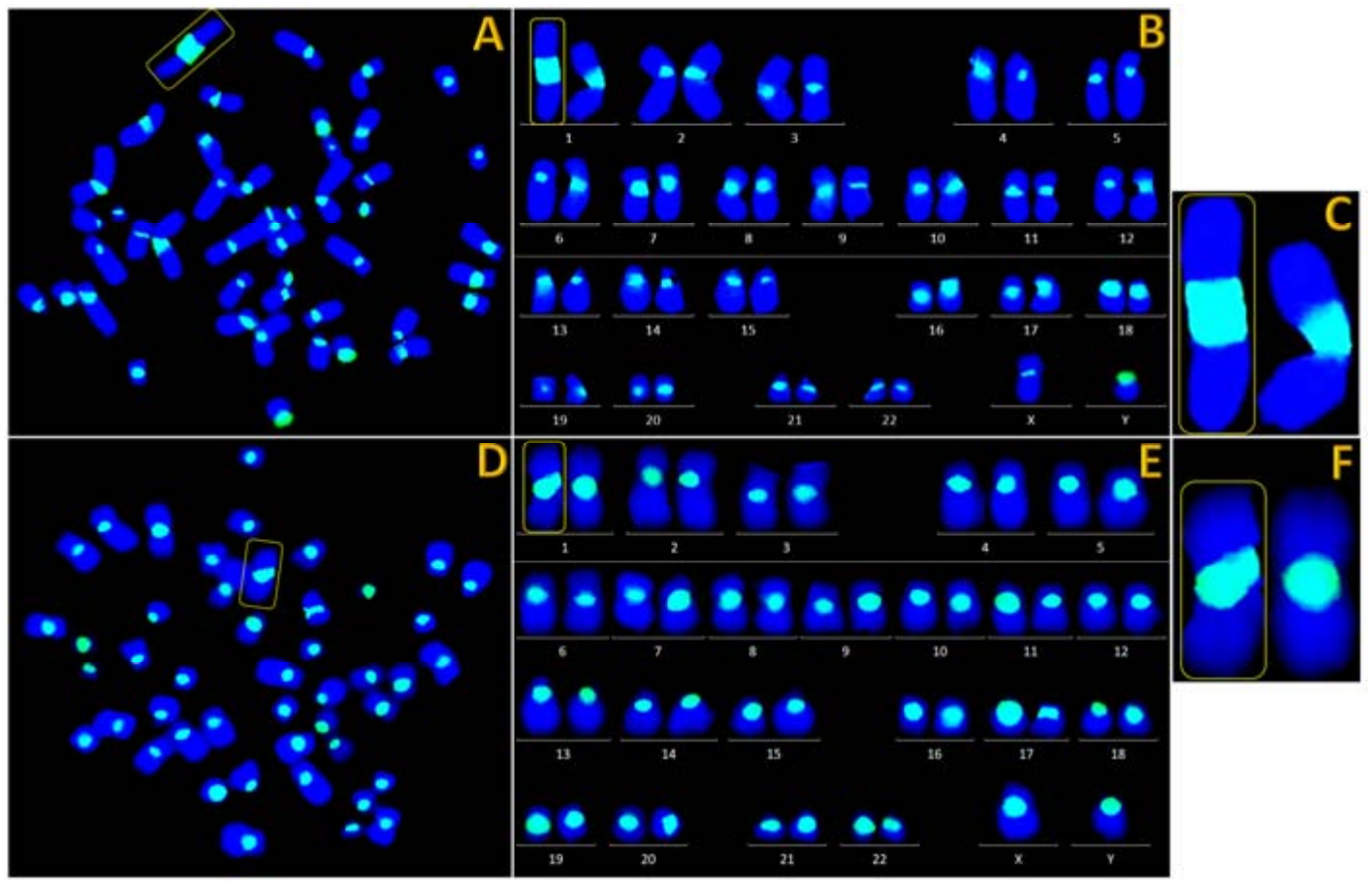
Representative metaphase chromosomes stained with pan-cent-FISH. (A-C) Metaphase from a subject exhibiting an enlarged centromere in one homolog of chromosome pair 1. (A) Representative metaphase spread. (B) Corresponding karyotype. (C) Enlarged view of chromosome pair 1 highlighting the enlarged centromere with increased centromeric signal (yellow box). (D-F) Representative metaphase from a healthy volunteer (control). (D) Metaphase spread. (E) Corresponding karyotype. (F) Enlarged view of chromosome pair 1 showing normal centromeric signal. Yellow boxes indicate the chromosome pair selected for detailed visualization.

**Figure 3:**
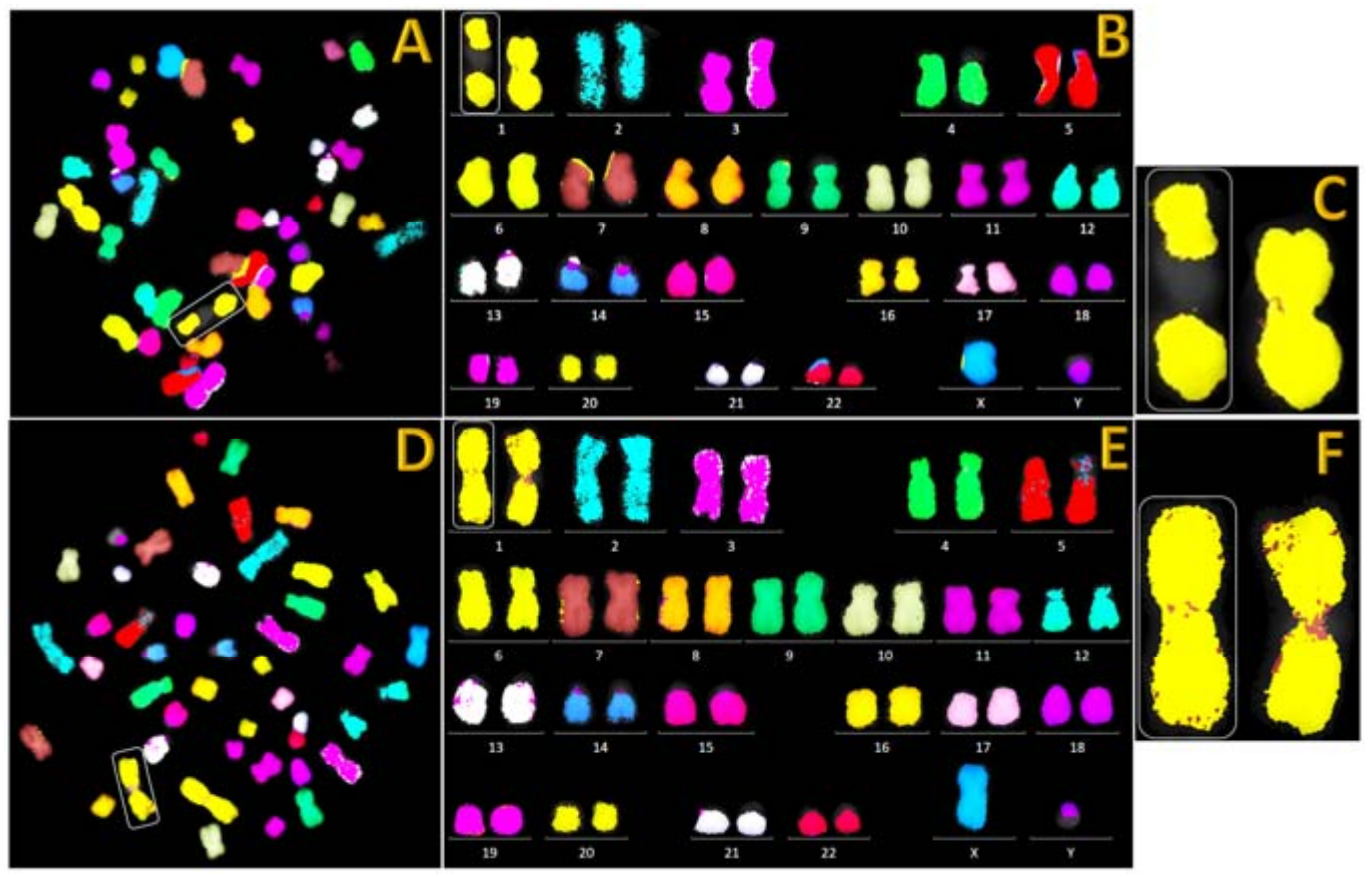
Representative metaphase chromosomes analyzed by mFISH. (A-C) Metaphase from a subject exhibiting an enlarged centromere in one homolog of chromosome pair 1. (A) Representative mFISH metaphase spread. (B) Corresponding karyotype. (C) Enlarged view of chromosome pair 1 highlighting the enlarged centromere (white box). (D-F) Representative mFISH metaphase from a healthy volunteer (control). (D) mFISH metaphase spread. (E) Corresponding karyotype. (F) Enlarged view of chromosome pair 1 showing normal chromosome morphology. White boxes indicate the chromosome pair selected for detailed visualization.

For quantitative assessment, the centromere-to-chromosome length ratio (CCR) was measured in pancent-FISH-processed metaphases, which provided optimal delineation of centromeric boundaries. Analysis of 50 metaphases per individual revealed a mean CCR of 0.314 ± 0.019 for the enlarged chromosome 1 homologue in the subject, compared with 0.169 ± 0.019 for the unaffected homologue of the same individual (Table 1). In the control, the mean CCR values for the first and second chromosome 1 homologues were 0.186 ± 0.038 and 0.161 ± 0.036, respectively, both in agreement with the unaffected homologue of the subject and with published normative values for chromosome 1. The enlarged homologue thus exhibited an approximately 1.86-fold increase in CCR relative to all chromosomes with normal centromeric morphology.

**Table 1.** Table shows the Centromere-to-Chromosome Ratio (CCR) of the 1st and 2nd homologues of Chromosome Pair 1 (CP1) for 50 cells each in the subject and control groups, with the mean and standard deviation given for each column.

| **S. No.** | **SUBJECT** |  | **CONTROL** |  |
|---|---|---|---|---|
|  | **CCR of 1<sup>st</sup> homologue of CP1** | **CCR of 2<sup>nd</sup> homologue of CP1** | **CCR of 1<sup>st</sup> homologue of CP1** | **CCR of 2<sup>nd</sup> homologue of CP1** |
| 1 | 0.307 | 0.196 | 0.239 | 0.169 |
| 2 | 0.307 | 0.169 | 0.222 | 0.159 |
| 3 | 0.316 | 0.179 | 0.233 | 0.178 |
| 4 | 0.310 | 0.184 | 0.152 | 0.114 |
| 5 | 0.315 | 0.205 | 0.239 | 0.222 |
| 6 | 0.321 | 0.208 | 0.167 | 0.149 |
| 7 | 0.274 | 0.147 | 0.217 | 0.200 |
| 8 | 0.252 | 0.182 | 0.250 | 0.232 |
| 9 | 0.305 | 0.155 | 0.242 | 0.203 |
| 10 | 0.301 | 0.211 | 0.130 | 0.104 |
| 11 | 0.317 | 0.157 | 0.259 | 0.196 |
| 12 | 0.322 | 0.160 | 0.128 | 0.117 |
| 13 | 0.321 | 0.206 | 0.164 | 0.149 |
| 14 | 0.321 | 0.148 | 0.228 | 0.212 |
| 15 | 0.304 | 0.186 | 0.190 | 0.177 |
| 16 | 0.319 | 0.167 | 0.174 | 0.167 |
| 17 | 0.321 | 0.183 | 0.137 | 0.118 |
| 18 | 0.308 | 0.163 | 0.188 | 0.171 |
| 19 | 0.312 | 0.146 | 0.169 | 0.158 |
| 20 | 0.329 | 0.169 | 0.208 | 0.152 |
| 21 | 0.316 | 0.156 | 0.133 | 0.119 |
| 22 | 0.323 | 0.155 | 0.135 | 0.091 |
| 23 | 0.310 | 0.160 | 0.167 | 0.152 |
| 24 | 0.308 | 0.154 | 0.125 | 0.087 |
| 25 | 0.286 | 0.145 | 0.163 | 0.158 |
| 26 | 0.313 | 0.143 | 0.113 | 0.100 |
| 27 | 0.323 | 0.151 | 0.164 | 0.153 |
| 28 | 0.316 | 0.193 | 0.250 | 0.238 |
| 29 | 0.324 | 0.210 | 0.216 | 0.216 |
| 30 | 0.319 | 0.160 | 0.204 | 0.154 |
| 31 | 0.313 | 0.161 | 0.192 | 0.182 |
| 32 | 0.316 | 0.169 | 0.196 | 0.146 |
| 33 | 0.304 | 0.172 | 0.200 | 0.167 |
| 34 | 0.375 | 0.184 | 0.145 | 0.167 |
| 35 | 0.321 | 0.182 | 0.160 | 0.160 |
| 36 | 0.330 | 0.159 | 0.200 | 0.207 |
| 37 | 0.312 | 0.158 | 0.238 | 0.214 |
| 38 | 0.333 | 0.156 | 0.200 | 0.167 |
| 39 | 0.317 | 0.161 | 0.211 | 0.151 |
| 40 | 0.321 | 0.150 | 0.144 | 0.125 |
| 41 | 0.294 | 0.149 | 0.167 | 0.112 |
| 42 | 0.304 | 0.137 | 0.179 | 0.148 |
| 43 | 0.323 | 0.154 | 0.200 | 0.180 |
| 44 | 0.297 | 0.182 | 0.196 | 0.173 |
| 45 | 0.308 | 0.161 | 0.213 | 0.170 |
| 46 | 0.280 | 0.162 | 0.180 | 0.161 |
| 47 | 0.330 | 0.176 | 0.150 | 0.135 |
| 48 | 0.361 | 0.171 | 0.200 | 0.120 |
| 49 | 0.346 | 0.172 | 0.154 | 0.164 |
| 50 | 0.300 | 0.163 | 0.171 | 0.162 |
| **Average** | **0.314** | **0.169** | **0.186** | **0.161** |
| **SD** | **0.019** | **0.019** | **0.038** | **0.036** |

Statistical comparison using a two-tailed Student’s unpaired t-test demonstrated that the CCR of the enlarged chromosome 1 homologue was significantly greater than that of all normal chromosome 1 homologues (t = 28.4, p < 0.0001). No statistically significant differences in CCR were observed among chromosomes with normal centromeric morphology across either individual. These findings provide quantitative evidence for a substantial and specific expansion of the centromeric region in a single chromosome 1 homologue of the subject.

## Discussion

This study reports the incidental identification of a constitutional centromeric heteromorphism of chromosome 1 in a phenotypically normal male referred for cytogenetic biodosimetry. Giemsa staining, pan-centromere FISH, and mFISH independently confirmed an ∼1.86-fold expansion of the centromeric region restricted to a single chromosome 1 homologue, with no dicentric chromosomes detected across 500 metaphases, excluding significant radiation exposure.

The consistent presence of the variant in all metaphases, unaffected contralateral homologue, and absence of phenotypic abnormality in the subject and first-degree relatives establish its constitutional, benign nature, consistent with published classification of chromosomal heteromorphisms as mitotically stable polymorphisms without clinical consequence (13, 14). Critically, the expanded pancentromere FISH signal at the primary constriction distinguishes this case from the more common 1qh+ variant, which involves enlargement of the pericentromeric 1q12 heterochromatic block rather than the centromere proper. Chromosome 1 harbours one of the largest and most polymorphic alpha satellite higher-order repeat (HOR) arrays in the human genome (15), and the observed centromeric expansion most likely reflects amplification or unequal recombination within this array, a mechanism consistent with the known recombinogenic susceptibility of chromosome 1 alphoid sequences (16). The absence of mitotic instability across all analysed metaphases confirms that this amplification does not compromise kinetochore function or chromosome segregation fidelity.

The quantitative CCR derived from pan-centromere FISH provides an objective metric rarely applied in heteromorphism case series, where qualitative banding assessment predominates. The highly significant CCR difference between the variant and all normal chromosome 1 homologues (t = 28.4, p < 0.0001) validates the CCR as a robust benchmark for centromeric size comparisons. mFISH, enabling simultaneous color discrimination of all 24 chromosomes (17), definitively excluded translocation or insertion of foreign material, confirming the expansion as intrinsic to chromosome 1. The convergence of three independent methods, each confirming the same variant homologue, represents a more rigorous characterization than is typically applied to constitutional centromeric variants.

An important practical implication of this finding relates to radiation biodosimetry and constitutional cytogenetics. Large centromeric heteromorphisms may be mistaken for structural abnormalities when identified during routine chromosome analysis. The combined application of pan-centromeric FISH, mFISH, and quantitative morphometric assessment enabled accurate characterization of the variant and exclusion of cryptic chromosomal rearrangements.

The study is limited by its single-case design and absence of molecular characterization of the expanded domain. Future work should apply long-read sequencing to resolve alpha satellite HOR copy number, use CENP-A chromatin immunoprecipitation to assess functional centromere competency, and undertake family studies to establish the inheritance pattern.

## Conclusion

A rare constitutional centromeric enlargement of chromosome 1 was identified and rigorously characterized in a phenotypically normal individual. The combined cytogenetic evidence confirmed a benign heteromorphic variant and excluded cryptic chromosomal rearrangements. This report broadens current knowledge of human centromeric variation and highlights the value of integrating molecular cytogenetic and quantitative approaches for accurate interpretation of atypical chromosome morphology in diagnostic and biodosimetric settings.

## Acknowledgements

The authors gratefully acknowledge the valuable assistance and technical support of Mr. Shrikant Jagtap and the technical staff of the Biodosimetry Laboratory, Radiological Physics and Advisory Division, Bhabha Atomic Research Centre.

## Statement of Ethics

### Study Approval Statement

This study was reviewed and approved by the Institutional Ethics Committee of the Bhabha Atomic Research Centre, Mumbai, India (Ref: BHMEC/NP/12/59/2026).

### Consent to Participate Statement

Written informed consent to participate was obtained from both participants (the subject and the healthy volunteer) before blood collection.

### Consent to Publish Statement

Written informed consent for publication of the details of the medical case and the accompanying images was obtained from the participants.

### Identifiability

The participants have been anonymized, and no identifying information is given in the text, tables or images.

### Author Contributions

K.B.S.: Investigation, methodology (cytogenetic and FISH analysis), data curation.

U.N.D.: Investigation, data curation, data curation.

A.N.: Investigation, formal analysis.

R.K.C.: Conceptualization, supervision, formal analysis, writing – original draft, writing – review and editing.

N.N.B.: Supervision, review and editing.

A.K.: Supervision, review and editing.

### Data Availability Statement

All data supporting the findings of this study are included in this article. Raw images and measurements are available from the corresponding author on reasonable request.

